# Evolution of Chi motifs in Proteobacteria

**DOI:** 10.1101/2020.08.13.249359

**Authors:** Angélique Buton, Louis-Marie Bobay

**Affiliations:** Department of Biology, University of North Carolina Greensboro, 321 McIver Street, PO Box 26170, Greensboro, NC 27402, USA

**Keywords:** homologous recombination, bacteria, genome evolution, DNA motifs

## Abstract

Homologous recombination is a key pathway found in nearly all bacterial taxa. The recombination complex allows bacteria to repair DNA double strand breaks but also promotes adaption through the exchange of DNA between cells. In Proteobacteria, this process is mediated by the RecBCD complex, which relies on the recognition of a DNA motif named Chi to initiate recombination. The Chi motif has been characterized in *Escherichia coli* and analogous sequences have been found in several other species from diverse families, suggesting that this mode of action is widespread across bacteria. However, the sequences of Chi-like motifs are known for only five bacterial species: *E. coli, Haemophilus influenzae*, *Bacillus subtilis*, *Lactococcus lactis* and *Staphylococcus aureus*. In this study we detected putative Chi motifs in a large dataset of Proteobacteria and we identified four additional motifs sharing high sequence similarity and similar properties to the Chi motif of *E. coli* in 85 species of Proteobacteria. Most Chi motifs were detected in *Enterobacteriaceae* and this motif appears well conserved in this family. However, we did not detect Chi motifs for the majority of Proteobacteria, suggesting that different motifs are used in these species. Altogether these results substantially expand our knowledge on the evolution of Chi motifs and on the recombination process in bacteria.

## Introduction

Bacteria frequently suffer DNA double-strand breaks; and these damages need to be efficiently repaired to avoid cell death. One of the primary repair mechanisms is mediated by homologous recombination which allows the exchange of homologous sequences and thus the repair of damaged DNA ^1^. Homologous recombination is also a key mechanism contributing to the rapid evolution of bacteria and their viruses ^2^. In Proteobacteria, the main enzymatic complex involved in this process is the RecBCD helicase-nuclease complex ^3,4^. Following a double-strand break, RecBCD binds to the broken end of the DNA and unwinds it ^5,6^ upon encountering a Chi (Crossover Hotspot Instigator) site. The recognition of the Chi motif triggers an endonucleolytic nick near the 3’ end of the Chi motif by RecBCD ^7^ and initiates the loading of RecA proteins on the DNA strand ^8^. The DNA-RecA complex can then initiate the search and exchange of genetic material with a homologous sequence ^7^. The recognition of Chi motifs, mediated by the RecC subunit ^9^, thus plays a key function in the initiation of the homologous recombination.

Chi sites have been well defined in *Escherichia coli*, where the sequence of this 8nt-long motif is 5’-GCTGGTGG-3’ ^10,11^. This motif is over-represented on the genome with one motif every five kilobases on average ^12^. Chi sites are also more frequent in the conserved regions of the bacterial genome (i.e. the core genome) than in the genes recently acquired ^13,14^ and in genomic repeats ^15^, suggesting the importance of the repair of the regions that encode the most essential cellular functions. This motif is also polarized on the chromosome, which is an important feature for the recognition of Chi sites by RecBCD ^16^: in *E. coli* Chi motifs are found almost exclusively on the leading strand of replication. Chi sites have an important role in the RecBCD pathway as its recognition by this enzymatic complex is a key step in initiating this mechanism. However, most of the knowledge on Chi sites has been derived from *E. coli* and little is known about this mechanism in other species.

Chi motifs and functionally-related motifs (i.e. Chi-like motifs) have only been identified in four other bacteria from diverse taxonomic groups: *Haemophilus influenzae* (5’-GNTGGTGG-3’/5’G(G/C)TGGAGG-3’)*, Bacillus subtilis* (5’-AGCGG-3’)*, Lactococcus lactis* (5’-GCGCGT-3’) ^17^ and *Staphylococcus aureus* (5’-GAAGCGG-3’) ^13^. Note that it remains unclear whether the 5’-AGCGG-3’ motif stimulates homologous recombination in *B. subtilis* and that it can therefore be considered a true Chi motif. For this reason, we will refer to “Chi motifs” to designate the motifs found in *E. coli* and related bacteria with a similar sequence. In contrast, we will use “Chi-like motifs” to designate all the motifs that are thought to be associated with homologous recombination in prokaryotes. For most of the species with Chi-like sequences identified, Chi-like sequences are statistically over-represented in the core genome ^17^. The presence of Chi-like motifs across highly divergent bacteria suggests that this mechanism of action is widespread in bacteria and possibly universal. Moreover, *H. influenzae* and *E. coli* are both Gammaproteobacteria and present related Chi sequences. Although Chi-like sequences have likely evolved toward different sequences in most species, this last case suggests that they are evolving relatively slowly and indicates that it might be possible to identify them with computational approaches.

In this study we searched for the presence of Chi motifs related to *E. coli*’s Chi motif across hundreds of species of Proteobacteria. By using sequence similarity, oligonucleotide frequency statistics and the known properties of Chi motifs (i.e. polarization on the leading strand and over-representation on the core genome) we detected the presence of candidate Chi motifs across 87 species of Proteobacteria. The identified motifs appear restricted to five sequences: (GCTGGTGG, GCTGGCGG, GCTGCTGG, GGTGGTGG and GCTGGAGG). We further used phylogenomic approaches to reconstruct the evolution of these sequences in Proteobacteria. Our results underline that these sequences are well conserved in *Enterobactericeae*, suggesting a common mode of action of homologous recombination in these species. However, we did not find related sequences across most Proteobacteria, indicating that Chi motifs are either absent in these lineages or possess highly divergent sequences.

## Materials and Methods

### Data collection

Genomic sequences were downloaded from GenBank in May 2018. One genome per species was selected. If multiple genomes were available, we selected the one with the most complete assembly (the one with the smallest number of contigs), and if a species presented multiple assemblies with the same quality, we randomly selected one of the genomes. Following this procedure, we obtained 8,924 genomes from distinct species. From this list, we extracted the species belonging to the Proteobacteria and the Terrabacteria, which yielded a total of 2603 species, composed of 1,071 and 992 species of Proteobacteria and Terrabacteria, respectively. We then selected the genomes that with complete assemblies (i.e. assemblies at the scaffold or the contig level were discarded), and this yielded a total or 495 Proteobacteria and 363 Terrabacteria. Dataset information such as species name, GenBank accession number, taxonomy, genome size and nucleotide composition (GC-content) are detailed in Dataset S1.

### Definition of the core genome

It has been previously observed that Chi sites are only statistically over-represented on the core genome ^17^, likely because these regions need to be repaired efficiently. In order to improve the sensitivity of our approach, we restricted our analyses of Chi motifs to the core genome of each species. The core genome of a bacterial species is typically defined by all the genes present in every or almost every strain of a species. However, it is not possible to build a core genome for most species, since most species present a single or a few sequenced genome(s). To circumvent this issue, we built a pseudo core genome for each species by using the set of orthologous genes shared among closely related species. Indeed, the core genes of a given species typically encode for the most essential cellular functions (typically housekeeping genes) and those are expected to be shared by closely related species. For each species, we used HMMer v3 ^18^ and the set of 43 universal protein profiles identified in a previous study ^19^. Each of the universal proteins identified were extracted and aligned with MAFFT v7 ^20^. For each alignment, poorly aligned regions were removed using Gblocks v0.91b ^21^. The 43 universal protein alignments were then concatenated into a single alignment. All pairwise distances were computed on the concatenate and the distances were used to identify all the pairs of most closely related species. For each pair of most related species, the set of shared orthologs was defined as best reciprocal hits using USEARCH v11 (global alignment) with a minimum of 70% sequence identity and 80% length conservation ^22^. For each species, the pseudo core genome was defined as the set of genes shared by the two most closely related species.

### Detection of Chi sites

Candidate Chi motifs were inferred by using multiple criteria. First, we searched for the presence of candidates Chi sites with an identical or highly similar sequence of the motif found in *E. coli* (5’-GCTGGTGG-3’). All the motifs with one nucleotide difference compared to the motif od *E. coli* have been analyzed, namely a total of 25 motifs: GCTGGTGG, ACTGGTGG, CCTGGTGG, TCTGGTGG, GATGGTGG, GGTGGTGG, GTTGGTGG, GCAGGTGG, GCGGGTGG, GCCGGTGG, GCTAGTGG, GCTTGTGG, GCTCGTGG, GCTGATGG, GCTGTTGG, GCTGCTGG, GCTGGAGG, GCTGGCGG, GCTGGGGG, GCTGGTAG, GCTGGTCG, GCTGGTTG, GCTGGTGA, GCTGGTGC, GCTGGTGT. The pseudo core genomes of the 2,063 species were analyzed by the software R’MES v3.1.0 ^23^. The statistics of all the possible octamers were performed using a Gaussian law ^24–26^ in R’MES v3.1.0, allowing the calculation of the expected frequency of the motifs relative to the oligonucleotide composition of the genomes. The over-representation of each motif was then evaluated by a statistical comparison between the observed and expected numbers using different Markov models to take into account the different levels of oligonucleotide composition of the sequences (different composition bias: mono-nucleotides, di-nucleotides, tri-nucleotides, quadra-nucleotides and penta-nucleotides and hexa-nucleotides). For each model, a motif was considered as significantly overrepresented based on the statistics observed for the Chi site frequencies of *E. coli* (thresholds used are presented Table S1). The detected motifs were considered as candidate Chi motifs when found significantly over-represented in the core genome with at least three different Markov models and when found polarized on the genome. The polarization statistics were computed with the Wilcoxon signed-rank test implemented in R version 3.5.1. Since we couldn’t confidently identify the origin and terminus of replication for all species, we reasoned that, in the absence of polarization on the genome, Chi sites should be randomly distributed on both DNA strands. If Chi sites are polarized on the chromosome, we should observe a biased distribution where consecutive Chi sites should be found on the same strand more frequently than expected by chance. We compared the distribution of the positions of Chi sites on the positive strand and on the negative strand. We considered that the candidate Chi motifs were polarized on the genome when the distribution was significantly biased (*p*-value < 0.001, Wilcoxon test) from the null expectation.

### Phylogenetic analysis

The concatenated alignment of 43 protein sequences described above was used to build the phylogenetic tree of the Proteobacteria ^19^. We built the phylogenetic tree with a maximum likelihood approach using RAxML v8 ^27^ with the LG model, which was selected as best-scoring model by RAxML. The tree was rooted by including the sequences of three Firmicutes: *Bacillus cereus*, *Lactobacillus casei* and *Staphylococcus saprophyticus*. The resulting tree was edited with colors and legends with iTOL ^28^. The Newick format of the tree is available in Dataset S2.

## Results

### Detection of five Chi motifs in Proteobacteria

We used multiple criteria to detect putative Chi sites in our dataset of 495 Proteobacteria: i) sequence similarity to the Chi motif of *E. coli*, ii) polarization with the leading strand of replication and iii) statistical overrepresentation on the core genome relative to the oligonucleotide composition of these genomes (see Methods for details). Because multiple oligonucleotide biases can impact the inference of Chi motifs, we computed the statistical overrepresentation of these motifs in the core genomes with R’MES using different Markov models: from model 0, which accounts for the mononucleotide composition of the core genome, to model 6, which accounts for the pentanucleotide composition of the core genome. A Chi motif was inferred as a candidate when statistically overrepresented in the core genome with three Markov models (see Table S2-3 for the detailed results across all Markov models). Using these different criteria, we identified 85 species of Proteobacteria with candidate Chi sites (see Dataset S3 for details). Across these species, five main motifs were identified as potential Chi motifs in Proteobacteria: GCTGGTGG, GCTGGCGG, GCTGCTGG, GGTGGTGG and GCTGGAGG (Figure 1). The most frequent motif, GCTGGTGG, is the Chi motif shown to be functional in *E. coli*.

**Figure 1:**
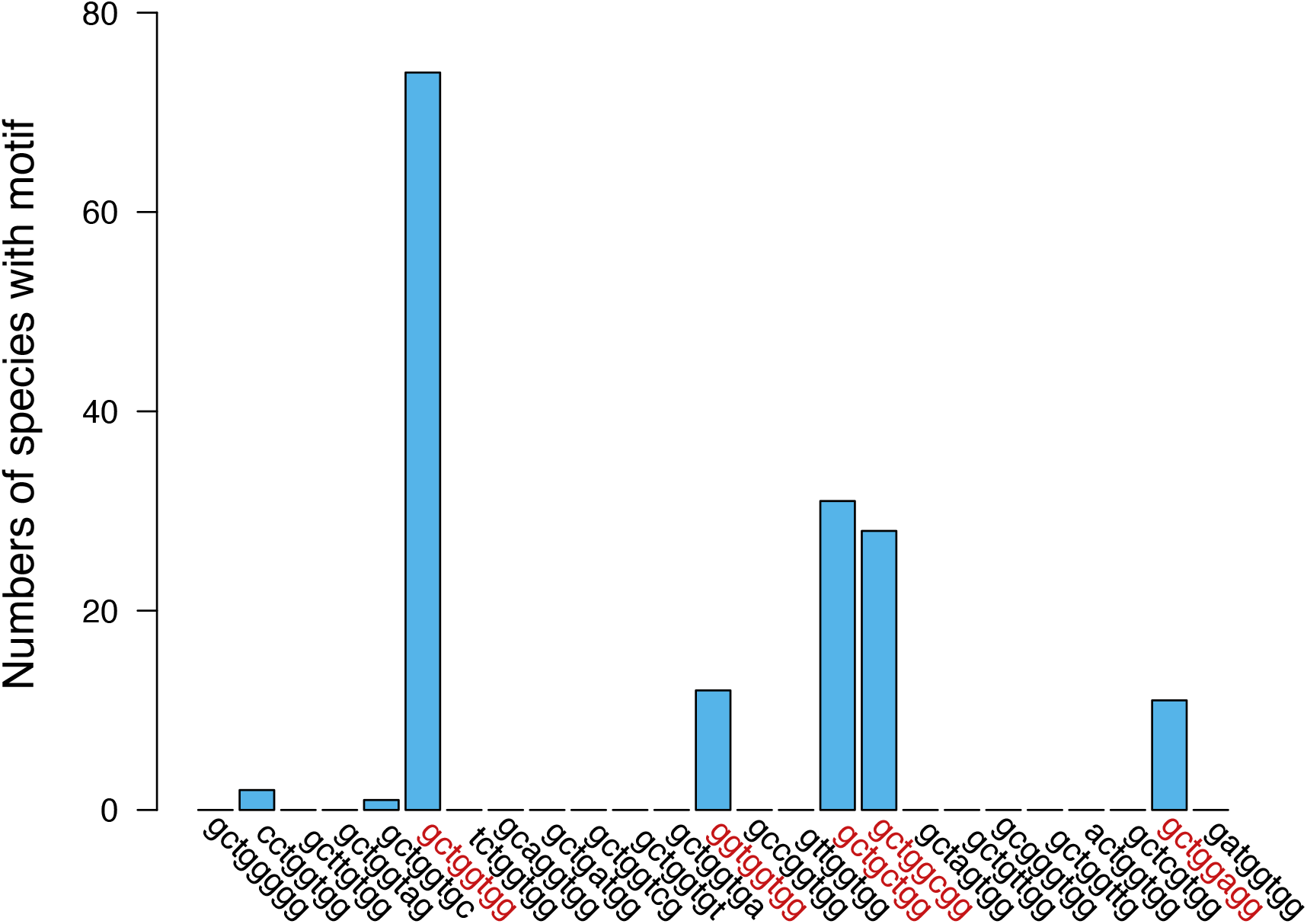
Frequencies and sequences of candidate Chi motifs identified in Proteobacteria. Five main Chi motifs have been detected in a total of 87 species of Proteobacteria based on our criteria.

### Estimation of false positives

To assess the quality of our detection approach, we used a large sample of Terrabacteria (n=363) to estimate our rate of false positive detection (see Table S1-3 for the detailed results across all Markov models). Terrabacteria are highly divergent from Proteobacteria but they also exhibit a large diversity of genome size and nucleotide composition. We reasoned that these divergent bacteria are not expected to present similar motifs to Proteobacteria. Indeed, the Chi motifs previously identified in *B. subtilis*, *L. lactis* and *S. aureus* are not related to the motifs inferred in Proteobacteria. Using the same detection procedure for the 25 putative motifs, we estimated our rate of false positives to be 3.9%. Interestingly, the putative Chi motifs are GC-rich and the false positives were mostly found in species of Terrabacteria with high GC-content, indicating that a weak detection bias due to sequence composition remains even though our framework should compensate for mononucleotide and oligonucleotide compositions.

### GC-content and Chi sites

The reference Chi motif of *E. coli* is a GC-rich sequence and all the related Chi motifs that we searched in these genomes present high GC-content (76.7% on average). We observed that the species with inferred candidate Chi motifs present higher genomic GC-content (60.8%) relative to the species without Chi motifs (respectively, 60.8% and 51.5%; *P*<10^−5^; Wilcoxon test; Figure S1A). This difference was also observed when limiting the comparison to Gammaproteobacteria (*P*<10^−5^; Wilcoxon test; Figure S1B). However, the genomic GC-content of Proteobacteria is much lower than the GC-content of Chi motifs (respectively, 60.8% and 76.6%; *P*<10^−5^; Wilcoxon test; Figure S1A). These results suggest that species harboring these Chi motifs tend to be relatively GC-rich, but their GC-content remains much lower than the GC-content of the motif itself.

### Evolution of Chi sites in Proteobacteria

We further analyzed the evolution of Chi motifs in Proteobacteria. To do so, we reconstructed the phylogenetic tree of Proteobacteria using a set 43 universal proteins previously identified ^19^. The distribution of species with inferred Chi motifs in the Proteobacteria tree confirms that the vast majority of these species are close relatives of *E. coli*: mostly *Enterobacteriaceae* (Gammaproteobacteria) (Figure 2). Several species of Gammaproteobacteria—other than the *Enterobacteriaceae*—were inferred to possess candidate Chi motifs (*Aeromonas*, *Zobellella*, *Pseudomonas* et *Kushneria*) but their genomes contain a GC-content of about 60%, which suggests that these could represent false positives. In the other groups of Proteobacteria, we detected Chi motifs among Betaproteobacteria (*Paraburkholderia* genus), as well as in Epsilon/Deltaproteobacteria (*Myxococcus* genus) and Alphaproteobacteria (*Azorhizobium caulinodan*s and *Komagataeibacter europaeu*s). Those species all present a relatively high GC-content (GC% > 60%), also suggesting that the Chi motifs inferred in these species likely represent false positive. For comparison, the GC-content of *E. coli* is 52.3%.

**Figure 2:**
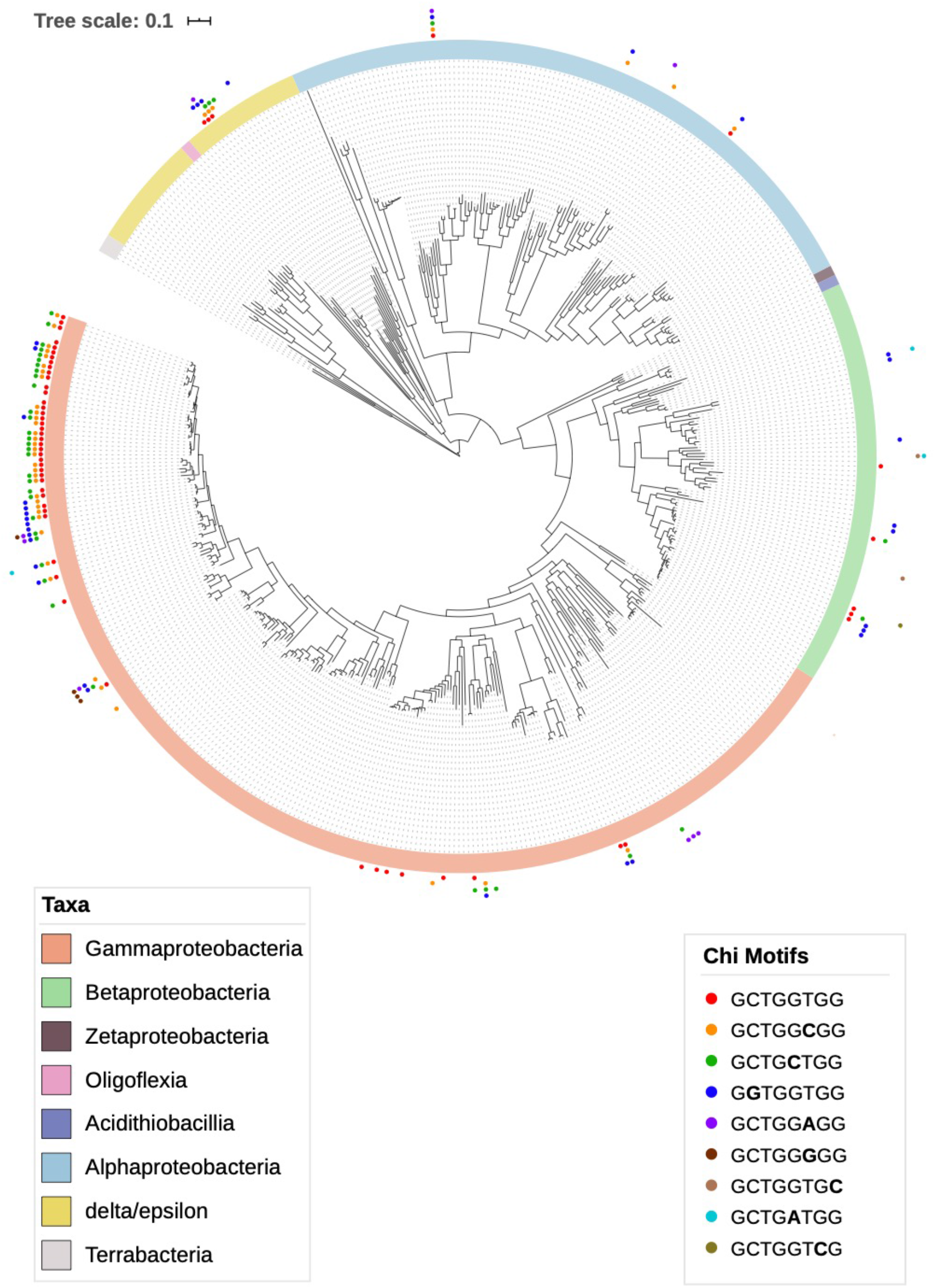
Phylogenetic tree of Proteobacteria and distribution of species with inferred Chi motifs. The tree was inferred using a set of 43 universal proteins and with RaxML (see Methods) using the LG substitution model and rooted using three Firmicutes species.

In contrast, virtually all *Enterobacteriaceae* presented Chi motifs. One notable exception is the genus *Yersinia* for which we did not detect the presence of Chi sites for any of the available species (Figure 3). The phylogenetic distribution of species with inferred Chi sites suggests that this motif was present in the ancestor of all *Enterobacteriaceae* and was subsequently lost in *Yersinia* and several *Serratia* species. Alternatively, it is possible that Chi sites were acquired after the divergence of the Yersinia/Serratia lineage and subsequently transferred to *Serratia proteamaculans*. Interestingly, many species of *Enterobacteriaceae* possess multiple Chi motifs. Indeed, three motifs are frequently identified in the same species: GCTGGTGG, GCTGGCGG and GCTGCTGG. This suggests that several species might recognize the degenerated Chi motif GCTG[G/C][T/C]GG. Finally, some of the more basally branching species of *Enterobacteriaceae* such as members of the genera *Dickeya* and *Edwardsiella* were predominantly found to present the motif GGTGGTGG. These results suggest, that Chi motifs are relatively well conserved across *Enterobacteriaceae* but that these sequences have experienced some minor sequence changes over time.

**Figure 3:**
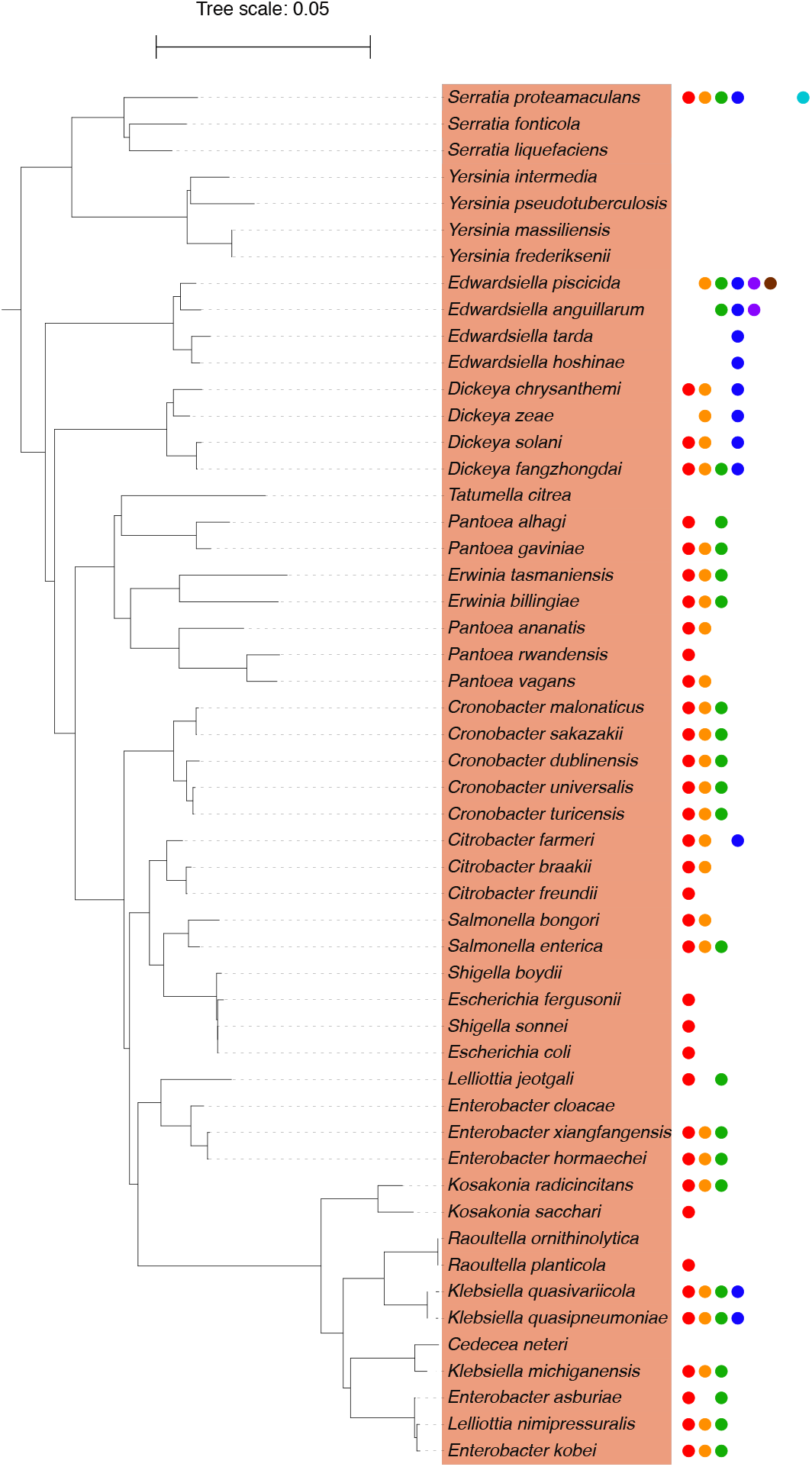
Distribution of candidate Chi motifs in *Enterobacteriacaea*. The subtree was extracted from the tree of Proteobacteria presented in Figure 2 and rooted with non-*Enterobacteriacaea* species.

### Extending the search of Chi motifs

Our initial analysis was based on stringent parameters to identify candidate Chi sites with high confidence using the same properties observed for *E. coli*’s Chi sites. It is very likely, however, that some Chi sites do not possess similar properties in other species. We conducted the same detection strategy without imposing any constraints on sequence polarity. This extended our list to a total of 234 species of Proteobacteria with candidate Chi motifs (Dataset S4) and these motifs were found in 64 Terrabacteria (21% of the species). Although this likely increases our rate of false positives, it is possible that many of these candidate motifs are true Chi motifs that need not be polarized to be functional.

We then extended the search of putative Chi motifs to more divergent sequences. Starting from the five candidate motifs that we detected, we generated 830 additional non-redundant motifs, each containing two different nucleotides relative to one of these five motifs. The same procedure was then used to search for statistically overrepresented motifs based on oligonucleotide compositions on the core genome and sequence polarity. We found very few polarized motifs statically overrepresented relative to oligonucleotide compositions of the genomes (Table S4). Using our previous criteria, we did not identify any of these motifs to be polarized and statistically overrepresented using at least three Markov models (Table S5). Finally, we searched for these 830 motifs without the polarity criterion and this yielded a list of only 11 species of Proteobacteria with candidate Chi motifs for 16 species of Terrabacteria (Dataset S5), indicating that these motifs are likely false positives. This result further suggests that reducing the stringency of our parameters does not substantially increase the list of species with candidate Chi motifs.

## Discussion

This study suggests that a relatively small proportion of Proteobacteria use Chi motifs. We found that a minority of Gammaproteobacteria (12.5% of the total Proteobacteria) is likely using this Chi motif. However, this study was limited to a list of motifs closely related to the one found in *E. coli*, i.e. a list of motifs with a one nucleotide difference (25 motifs). Among the 25 sequences, only five were found frequently in the Proteobacteria: 5’-GCTGGTGG-3’, 5’-GCTGGCGG-3’, 5’-GCTGCTGG-3’, 5’-GGTGGTGG-3’ and 5’-GCTGGAGG-3’. it is possible that some of these species of Proteobacteria do not have Chi sites. Indeed, some of these species completely lack (e.g. *Buchnera aphidicola*) or show a strong reduction in homologous recombination (e.g. *Yersinia pestis*) ^29^. In addition, some species of Proteobacteria, such as Alphaproteobacteria, rely on the AddBC recombination system instead of RecBCD ^4^. However, given the focus of our procedure on Chi sites related to *E. coli*’s motif, it is very likely that more divergent—or unrelated—Chi-like sequences exist in Proteobacteria. Searching for additional motifs could enable the identification of Chi sites in these species. This endeavor remains difficult due the sensitivity of *ab initio* detections and that no Chi-like motifs unrelated to *E. coli*’s motif have been identified in Proteobacteria so far. In addition, it may not be possible at all to detect such motifs using computational approaches if these sequences are simply not statistically over-represented in the genome (see below).

Given the large dataset used for this study, it is likely that some Chi motifs identified correspond to detection errors (i.e. false positives). By using a large set of Terrabacteria species, which are thought to use different motifs ^17^, we identified our rate of false positives to reach about 4% of detected motifs. The detection of false positives seems to be closely linked to the richness in GC-content since most false positives were found in GC-rich genomes (e.g. Actinobacteria). Although the nucleotide composition biases were taken into account by the statistics generated by R’MES, it seems that these filters did not completely eliminate all the false positives due to nucleotide composition. This suggests that the candidate Chi motifs inferred in GC-rich species should be taken with care.

The evolution of Chi sites and other DNA motifs is still poorly understood; however, we can conclude that species related to *E. coli* likely use very similar motifs, indicating that these motifs have been conserved for millions of years. Based on our phylogenetic analysis we can infer that this motif appeared at least since the divergence between the *Enterobacteriaceae* and the rest of the Gammaproteobacteria and that the sequence of the motif evolved in some of these taxa. In agreement with our results, it was previously observed that several species of *Enterobacteriaceae* were capable of Chi-dependent cleavage of DNA constructs containing *E. coli*’s Chi motif ^30,31^. Our results extend these findings by showing that most *Enterobacteriaceae* contain polarized and statistically overrepresented Chi motifs in their core genomes and that some of these species likely use slightly different motifs than *E. coli*’s motif (or a degenerated version of this motif). However, we found that the majority—but not all— species of *Enterobacteriaceae* use Chi motifs similar to *E. coli*’s motif. Indeed, we did not detect the presence of polarized Chi motifs in *Vibrionaceae*, although several of these species were shown capable of Chi-dependent cleavage of DNA constructs containing *E. coli*’s Chi motif ^30^. However, we identified a non-polarized candidate Chi motif (GCTGGTGG) in 23 species of *Vibrio* (Dataset S4), indicating that polarity might not be required for the proper function of Chi motifs in these species. We did not detect candidate Chi motifs in *H. influenzae* (*Pasteurellaceae*), although Chi motifs were identified for this species in a previous study ^17^. In fact, the original study that identified Chi sites in *H. influenzae* ^17^ showed that these motifs are degenerated, less represented statistically and less polarized than the ones found in *E. coli*, even though their function has been experimentally validated ^32^. We used stringent criteria to avoid the detection of false positives, but it is possible that some species present Chi motifs too degenerated to be confidently identified using *in silico* approaches. Furthermore, it is likely that other species of Proteobacteria contain Chi sites with other sequences than those we searched for in this study. It is possible that the stringency of our approach prevented us from identifying additional motifs but relaxing our search criteria directly leads to a much higher rate of false positives. Our results do not constitute a proof that most Proteobacteria are devoid of Chi or Chi-like motifs, but our analyses rather suggest that their putative motifs must substantially differ from *E. coli*’s motif. Considering that some experimentally characterized motifs, such as the ones of *H. influenzae*, virtually escape to all bioinformatic detection procedure, it is possible that only an experimental approach would allow to confidently detect new Chi-like motifs in Proteobacteria. Together, these results contribute to better characterizing the signals and pathways involved in homologous recombination in bacteria, as well as the evolution of genomic motifs.

Overall, our results indicate that Chi motifs similar to *E. coli*’s motif evolved at least since the ancestor of *Enterobacteriaceae* and were lost in several lineages. Some of these species likely use motifs whose sequence is slightly different from *E. coli*’s motif. Using *in silico* approaches we did not confidently detect such motifs in other species of Proteobacteria, suggesting that these lineages use unrelated sequences or that these motifs do not present all the characteristics of *E. coli*’s motif (i.e. polarization on the leading strand). Experimental analyses will be needed to confirm that these motifs can efficiently trigger RecBCD-mediated homologous recombination. Overall this study revealed new candidate Chi motifs and extends our understanding of the mechanisms controlling homologous recombination in Proteobacteria.

## Acknowledgments

We thank Shealynn Burgess for assistance with the analyses, Kasie Raymann for criticisms on the manuscript and Caroline Stott for assistance with the figures. This work was supported by the National Science Foundation under Grant No. DEB-1831730 by the National Institute Of General Medical Sciences of the National Institutes of Health under Award Number R01GM132137.

## Authors’ contributions

LMB and AB designed the study. LMB and AB analyzed the data. LMB and AB wrote the manuscript. All authors read and approved the final manuscript.

---

**Figure S1:**
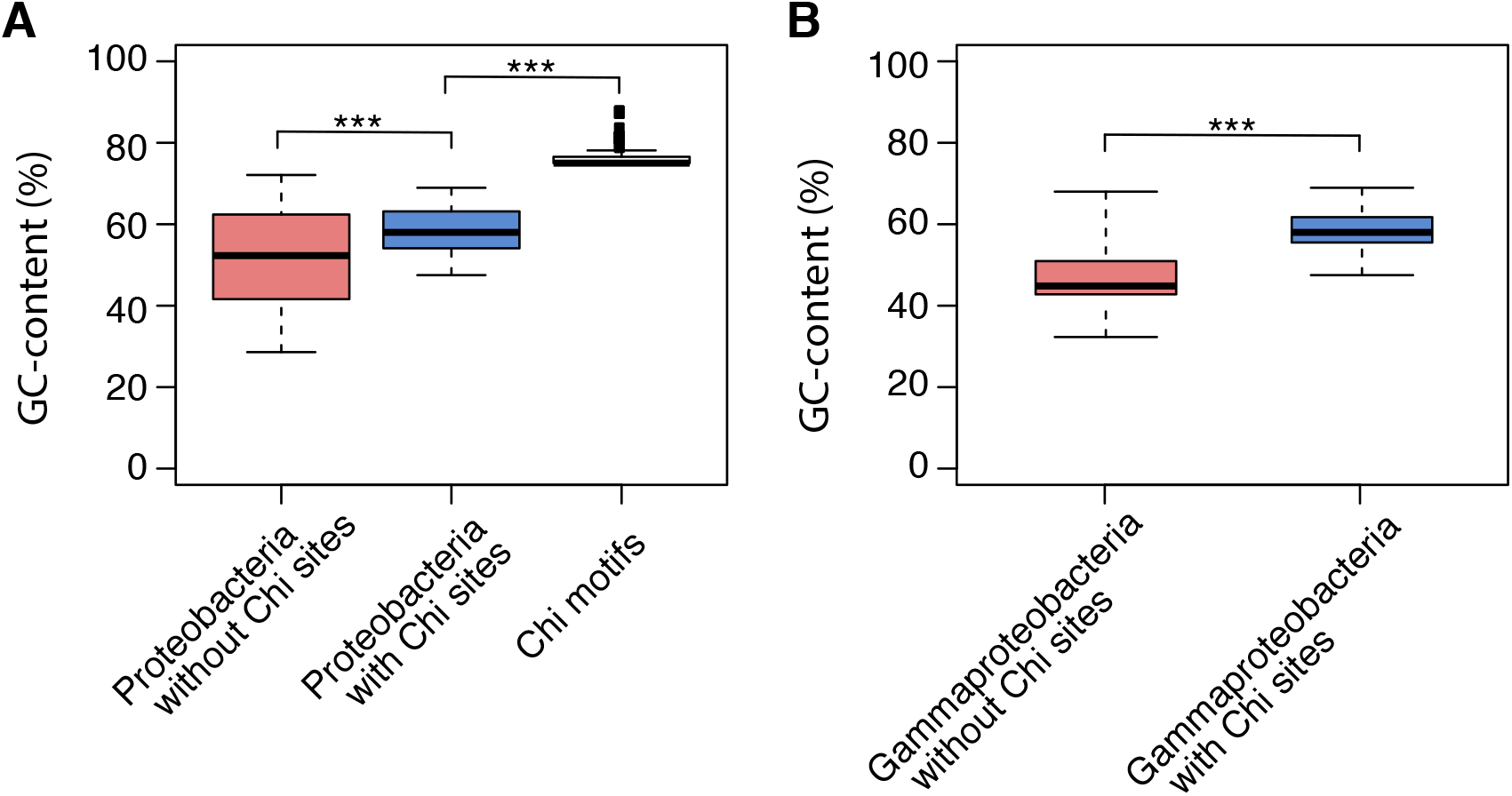
GC-content of Proteobacteria with or without inferred Chi sites. **A** comparison of GC-content between species of Proteobacteria with inferred Chi sites and without inferred Chi sites. The GC-content of the candidate Chi motifs is indicated in the third boxplot. **B** comparison of GC-content between species of Gammaproteobacteria with inferred Chi sites and without inferred Chi sites. (***) *P*<10^−5^, Wilcoxon test.

## Supplementary Tables and Datasets

**Table S1.** Statistical overrepresentation of Chi motifs across Markov models.

**Table S2.** Number of species identified with Chi motifs across Markov models (25 motifs searched).

**Table S3**. Number of species identified with Chi motifs 25 motifs searched).

**Table S4.** Number of species identified with Chi motifs across Markov models (830 motifs searched).

**Table S5**. Number of species identified with Chi motifs (830 motifs searched).

**Dataset S1.** List of analyzed genomes and genomic features.

**Dataset S2**. Phylogenetic tree of Proteobacteria in Newick format.

**Dataset S3.** List of species with candidate Chi motifs (25 polarized motifs searched).

**Dataset S4.** List of species with candidate Chi motifs (25 non-polarized motifs searched).

**Dataset S5.** List of species with candidate Chi motifs (830 non-polarized motifs searched).

